# Whole-Process 3D ECM-Encapsulated Organoid-Based Automated High-Throughput Screening Platform Accelerates Drug Discovery for Rare Diseases

**DOI:** 10.1101/2024.10.08.617181

**Authors:** Zhaoting Xu, Hui Yang, Yuru Zhou, Emmanuel Enoch Dzakah, Bing Zhao

## Abstract

The use of organoids, especially patient-derived organoids, for high-throughput screening (HTS) is widely accepted due to their ability to mimic the three-dimensional (3D) structure, function, and drug responses of in vivo tissues. However, the complexity of handling extracellular matrix (ECM) components with traditional HTS devices leads to the utilization of suspension cultures in matrix-free or matrix-low conditions during HTS, which can alter their transcriptomic landscape and drug responses. Here, we develop a whole-process 3D ECM-encapsulated organoid-based automated HTS (wp3D-OAHTS) platform, which enables the rapid and accurate generation of uniformly distributed 3D cell-matrix mixture domes at the center of each well in 96-well plates. This approach replicates the process of manual organoid culture but with superior stability and reproducibility. Utilizing this platform, we screened 2,802 compounds on neuroendocrine cervical cancer organoids, a rare malignancy with significant unmet clinical needs. We identified 7 top hits that display strong anti-tumor effects with remarkably low half-maximal inhibitory concentration (IC_50_) and validated the in vivo efficacy of Quisinostat 2HCl. Additionally, we demonstrated that employing 3D ECM-encapsulated organoid cultures for HTS, rather than suspended cultures, provides optimal conditions for drug discovery. Our wp3D-OAHTS platform significantly improves the rapidity and efficiency of new drug discovery for rare diseases.

## Introduction

The application of organoids for ex vivo high-throughput screening (HTS), particularly patient-derived organoids (PDOs), has gained widespread acceptance in drug development and precision medicine^1–5^. HTS allows for the rapid assessment of the sensitivity and resistance of thousands of drugs based on their cellular responses, independent of a full understanding of the molecular vulnerabilities underlying specific diseases. This process is advantageous in enabling the quick discovery of treatments for rare diseases lacking genomic mutation information^6^. Compared to conventional two-dimensional (2D) cell lines, organoids with a three-dimensional (3D) structure are more complex, closer to the architecture and behavior of native tissues, and more accurate in predicting drug responses^7^. Compared to patient-derived xenograft (PDX) models, PDOs eliminate of the confounding variables associated with interspecies differences and are more cost-effective, time-saving, and resource-efficient.

Organoids are typically developed from stem cells or progenitor cells encapsulated in extracellular matrix (ECM), which is crucial for the establishment of tissue-like architectures, cell-cell communications, and cell-ECM interactions^8–10^. However, previous attempts to employ organoids for HTS have frequently utilized matrix-free or matrix-low suspension culture conditions^1, 3, 11^, due to the challenges encountered by traditional automated HTS platforms when handling 3D ECM-encapsulated organoids. On one hand, the viscosity of most extracellular matrix components is temperature-dependent, meaning that cell-matrix mixtures will rapidly solidify within the pipette tips when exposed to room temperature during handling by traditional liquid dispensers, thus hindering the dispensing process^12, 13^. On the other hand, the requirement for cell-matrix mixtures to be centrally located in wells poses a significant obstacle when using high-density microplates, such as 96-well or 384-well plates, where well dimensions are exceedingly small. This small size dramatically increases the complexity of automated seeding of the ECM-encapsulated organoids. Therefore, organoid cultures were twisted to adapt organoid screenings to the conventional liquid dispenser-based HTS platforms, where the cells are suspended in matrix-low conditions rather than encapsulated in 3D ECM. However, even short-term suspension of organoids can lead to changes in their transcriptomic profile (Supplementary Fig. 1b, c) including the expression of tumor markers (Supplementary Fig. 1d) and significant modifications in key signaling pathways (Supplementary Fig. 1e), which may affect drug sensitivity among tumor cells.

To ensure that organoids retain maximal fidelity to in vivo tissues and to enhance the efficacy of drug screenings, we developed a flexible automated 3D ECM-encapsulated organoids spotter that enables the rapid and accurate generation of uniformly distributed 3D cell-matrix mixture domes at the center of each well in 96-well plates. By integrating a robotic arm, we combined our 3D ECM-encapsulated organoids spotter with three critical modules (liquid dispensing module, incubators, and detector) into an integrated workflow, realizing the establishment of a whole-process 3D ECM-encapsulated organoid-based automated HTS (wp3D-OAHTS) platform. Employing this platform, we conducted a high-throughput screen of 2,802 small molecules against neuroendocrine cervical cancer (NECC) organoids and identified seven compounds with half-maximal inhibitory concentration (IC_50_) in the low nanomolar range. The in vivo efficacy of a representative candidate Quisinostat 2HCI, which has not yet been approved by the FDA, was validated, suggesting a more effective suppression of the growth of NECC xenografts compared to drugs currently used in the clinic. Moreover, we found that our wp3D-OAHTS platform allows for more precise drug responses as compared to organoids in suspension. This proof-of-concept screening demonstrated the potential of our wp3D-OAHTS platform for anticancer drug discovery, especially for rare tumors.

## Results

### Workflow design and system development of a wp3D-OAHTS platform to innovate drug discovery

The wp3D-OAHTS platform we developed consists of four principal modules: a 3D ECM-encapsulated organoids spotter, a liquid dispensing module for culture medium addition and drug administration, an automated incubating module for organoid culture, and a detecting module for organoid imaging and cell viability testing (Fig. 1a, b). Among them, the 3D ECM-encapsulated organoids spotter is the key of handling ECM components during organoid screenings. Important design features of this spotter include a sample plate equipped with a cooling unit for storing and keeping cell-matrix mixtures at low temperatures before aspirating and spotting. To prevent cell sedimentation in the ECM, the mixtures in the sample plate will be automatically mixed thoroughly before dispensing. Pressure sources are used to control the flow rate of cell-matrix mixtures, and the spotting volume was proportional to and correlated with the plateau time. The homogenized cell-matrix mixtures are then aspirated by an 8-channel ECM spotting head with disposable tips and directly spotted into the wells (3 μl per well) in flying mode, with each well requiring only 0.1 seconds (Fig. 1c, Supplementary Video 1). Subsequently, the spotted plates undergo gelation on a heating unit until the ECM solidifies completely. A 96-channel liquid dispensing head is then used to add medium into all wells of the plate at once, taking just 0.07 seconds per well. After this, the plates are transferred by a robotic arm to incubators for culture. On a daily standardized schedule, the robotic arm transfers the plates from the incubators to the detector for data collection (e.g., imaging), a process that is accomplished in 3 seconds per well, providing automated continuous monitoring of organoid growth (Supplementary Video 2). On the day of drug administration, the robotic arm retrieves the plates from the incubator and transports them to the liquid dispensing module for medium replacement and drug administration, averaging 2 seconds per well, enabling automated large-scale drug dispensing (Supplementary Video 3). After 5-7 days of drug treatment, plates are transferred to the liquid dispensing module for medium removal and cell viability assay reagents addition, and then transferred to the incubators for incubation, requiring a scant 2 seconds per well. Finally, the plates are transferred to the detector for cell viability testing, with each well analyzed in a mere 1 second.

**Figure 1.**
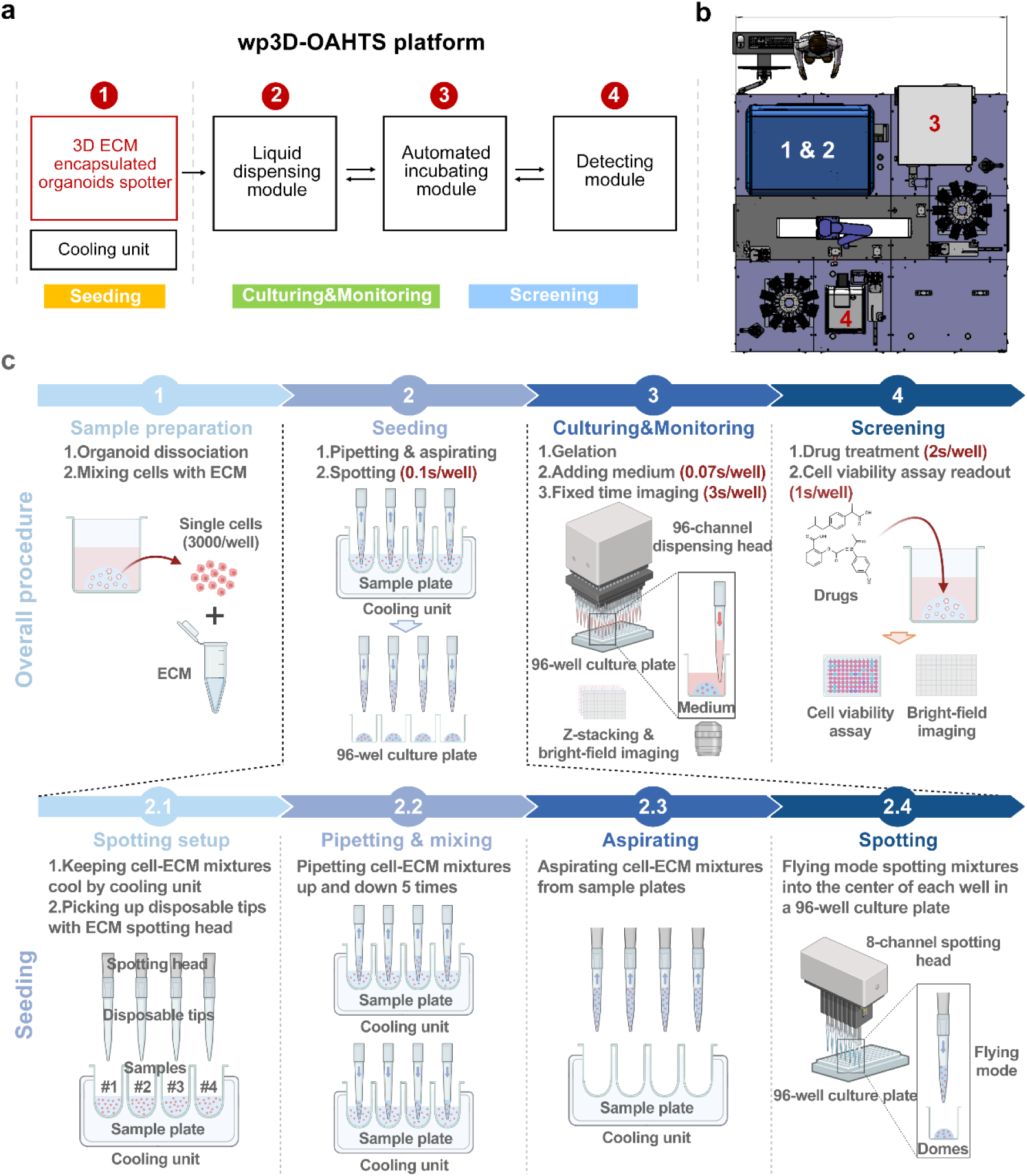
∣ Whole-process 3D ECM-encapsulated organoid-based automated high-throughput screening (wp3D-OAHTS) platform. (**a**) Four key modules of the wp3D-OAHTS platform. (**b**) The top view design of the wp3D-OAHTS platform. The number of each module in (a) was annotated on the corresponding device. (**c**) Top: Illustration of the overall procedure for conducting 3D organoid drug screening using the wp3D-OAHTS platform. Bottom: Detailed presentation of the most innovative automated seeding process establishing uniformly distributed 3D ECM-encapsulated organoid domes. The image was created with https://www.biorender.com/ (accessed on September 29, 2024).

In summary, when operating at full capacity, the wp3D-OAHTS platform can execute the spotting, data collection, drug treatment, and detection across 26,083 wells within a 10-day screening cycle including 5 days of culturing post-seeding and 5 days of drug treatment. When testing three replicates per drug, we could handle 8,694 different drugs within the same cycle. The efficiency and throughput of the platform mark a significant advancement in 3D ECM-encapsulated organoid-based high-throughput screening capabilities, demonstrating its potential for expediting drug discovery and personalized medicine.

### The automated platform greatly improves the homogeneity and reproducibility of 3D ECM-encapsulated organoid cultures compared to manual operation

The homogeneity and reproducibility of 3D ECM-encapsulated organoid cultures are critical prerequisites for performing 3D organoid-based HTS. To validate the superior consistency of our wp3D-OAHTS system in generating uniform ECM domes, we first compared its performance against manual seeding using cell-free ECM. Operator-to-operator variability and batch-to-batch inconsistencies are prominent drawbacks of manual seeding. In contrast, our automated spotter ensures precise central positioning within each well (Fig. 2a, d). Moreover, in a fully automated 96-well plate, the size and shape of the ECM domes exhibited minimal inter-well variations (Fig. 2b, c, e, f), establishing a solid foundation for the generation of stable and homogeneous 3D ECM-encapsulated organoids.

**Figure 2.**
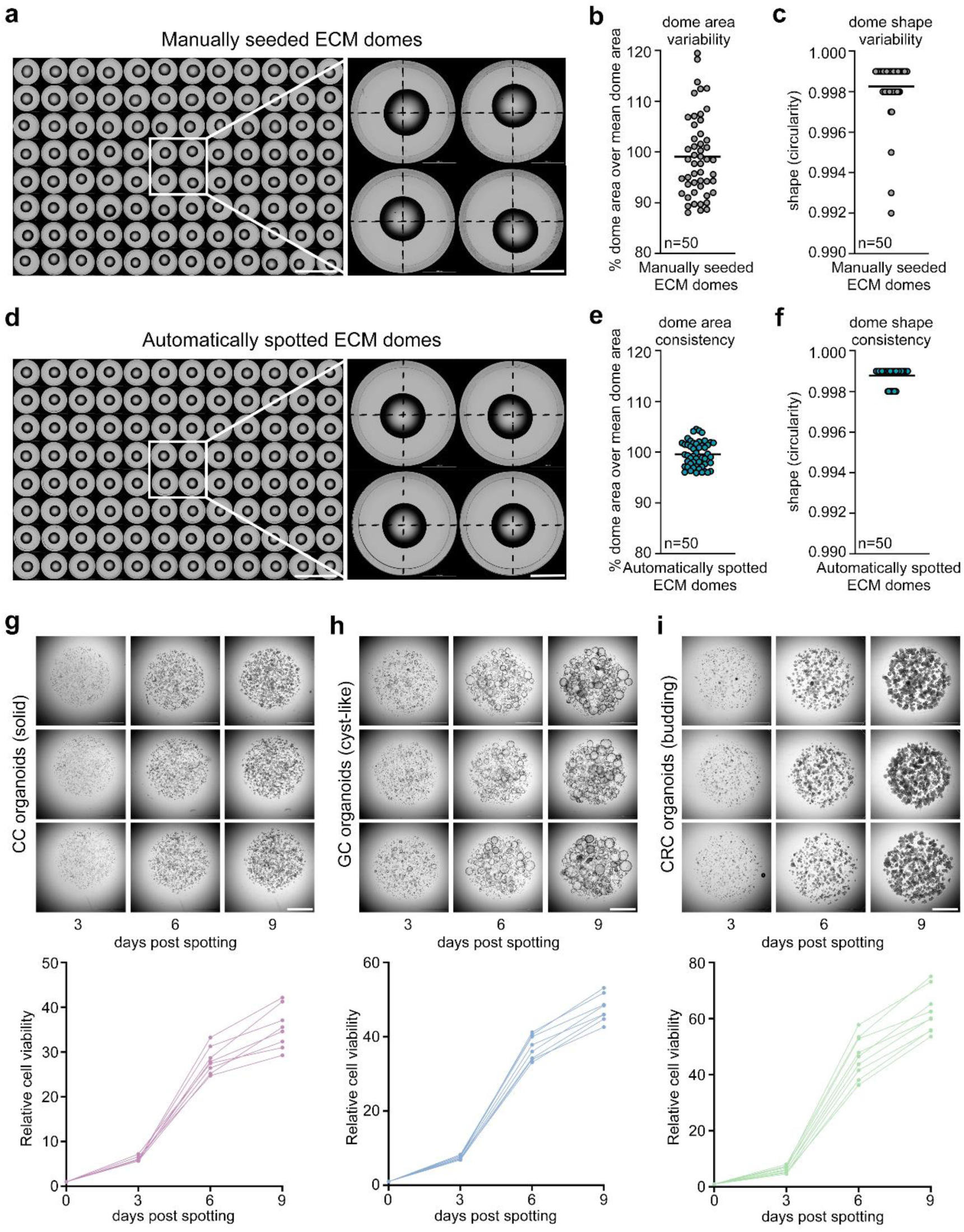
∣ The automated platform greatly improves the homogeneity and reproducibility of 3D ECM-encapsulated organoid cultures compared to manual operation. (**a**) Overview of an entire 96-well plate filled with manually seeded ECM domes (left) and enlarged view (right). Scale bars, 10 mm (left) and 2 mm (right). (**b**) Quantification of the dome-to-dome area variability of manually seeded ECM domes (dots). (**c**) Quantification of the shape (circularity) of manually seeded ECM domes (dots). (**d**) Overview of an entire 96-well plate filled with automatically spotted ECM domes (left) and enlarged view (right). Scale bars, 10 mm (left) and 2 mm (right). (**e**) Quantification of the dome-to-dome area variability of automatically spotted ECM domes (dots). (**f**) Quantification of the shape (circularity) of automatically spotted ECM domes (dots). (**g-i**) Top: Representative bright-field images showing cervical cancer (g), gastric cancer (h), and colorectal cancer (i) organoids growing after being automatically spotted. Scale bar, 1 mm. Bottom: Quantification of the cell viability change ratio of cervical cancer (g), gastric cancer (h), and colorectal cancer (i) organoids (dots) after being automatically spotted.

To ensure that the spotting process in the wp3D-OAHTS platform did not impair the biological activity of organoids, we mixed the dissociated single cells derived from PDOs of cervical cancer (CC), gastric cancer (GC), and colorectal cancer (CRC) respectively with ECM. The cell-matrix mixture domes (3,000 cells per dome) were generated in centers of wells in 96-well plates using the 3D ECM-encapsulated organoids spotter, and the growth of these organoids was continuously tracked via the automated monitoring pipeline in wp3D-OAHTS platform. These organoids were selected due to their distinct morphological characteristics such as solid, cyst-like, and budding structures. All of the organoids seeded by our spotter exhibited robust proliferation and rapid growth over time, with consistent development maintained across different wells throughout the culture (Fig. 2g, h, i). These results validated the capability of our platform in generating uniform, reproducible 3D ECM-encapsulated organoids that retain robust biological activity, positioning it as a high-precision, reliable system for large-scale 3D organoid-based HTS.

### The wp3D-OAHTS platform enables 2,802 drug screening on human neuroendocrine cervical cancer (NECC) organoids

The next aim is to demonstrate the potential of the platform for high-throughput automated 3D ECM-encapsulated organoid-based drug screening. To perform a proof-of-concept 3D organoids HTS for drug discovery in rare diseases, we focused on human neuroendocrine cervical cancer (NECC) organoids derived from a patient diagnosed with NECC and screened 2,802 small molecules using this organoid line. NECC is classified within the broader category of neuroendocrine neoplasms (NENs), which comprise well-differentiated neuroendocrine tumors (NETs) and poorly differentiated neuroendocrine carcinomas (NECs)^14–17^. Patients with NECs typically have a median life expectancy of less than one year^18^, whereas those diagnosed with NETs exhibit slightly improved outcomes, albeit with high and unpredictable variations in survival rates^19^. NENs have the highest incidence in the gastroenteropancreatic (GEP) system and lung, but they can also occur in other organs. For example, they are referred to as NECC when developing in the cervix. NECC is an aggressive histological subtype of cervical cancer accounting for about 1-1.5% of all cervical cancers, with small cell NECs being the most common form^20, 21^. Due to the rarity of this malignancy, there are few prospective studies or randomized trials specific to NECC to guide the standard of care management and therapy. Consequently, the clinical treatment protocols for patients with NECC primarily derive from those established for general cervical cancer or pulmonary neuroendocrine tumors^14^. Therefore, there is an urgent need to develop novel therapeutic agents and provide innovative treatment guidance for patients suffering from NECC.

To identify potential inhibitors for NECC, we obtained a tumor specimen from the surgical resection of an NECC patient and established the corresponding organoid line. The organoids demonstrated rapid proliferation in vitro with the typical solid structure (Fig. 3b, c). NECC organoids were dissociated, mixed with ECM, and automatically spotted into 129 96-well plates in 3D ECM-encapsulated conditions using our wp3D-OAHTS platform. 5 days later, 2,802 compounds (10μM) from libraries containing signaling pathway inhibitors, active pharmaceutical ingredients (APIs), natural products, and chemotherapeutic agents, some of which have been approved by the Food and Drug Administration (FDA), were added to the plates. After an additional 5 days of drug treatment, cell viability was assessed using a LivingCell-Fluo™ Organoid Vitality Assay (Fig. 3a). A total of 583 compounds with inhibition rates of more than 90% were chosen as the primary hit compounds (Fig. 3d). The *Z’*-factors were above 0.7 and signal-to-background (S/B) ratio parameters exceeded 8 in all screening plates (Fig. 3e, f), indicating a robust and high-quality HTS performance. The results from two independent assays of 77 randomly selected compounds revealed a high interexperiment correlation (Pearson’s coefficient of 0.86, *P* < 0.0001) (Fig. 3g). These results collectively demonstrated the accuracy, robustness, and reproducibility of our drug screening platform.

**Figure 3.**
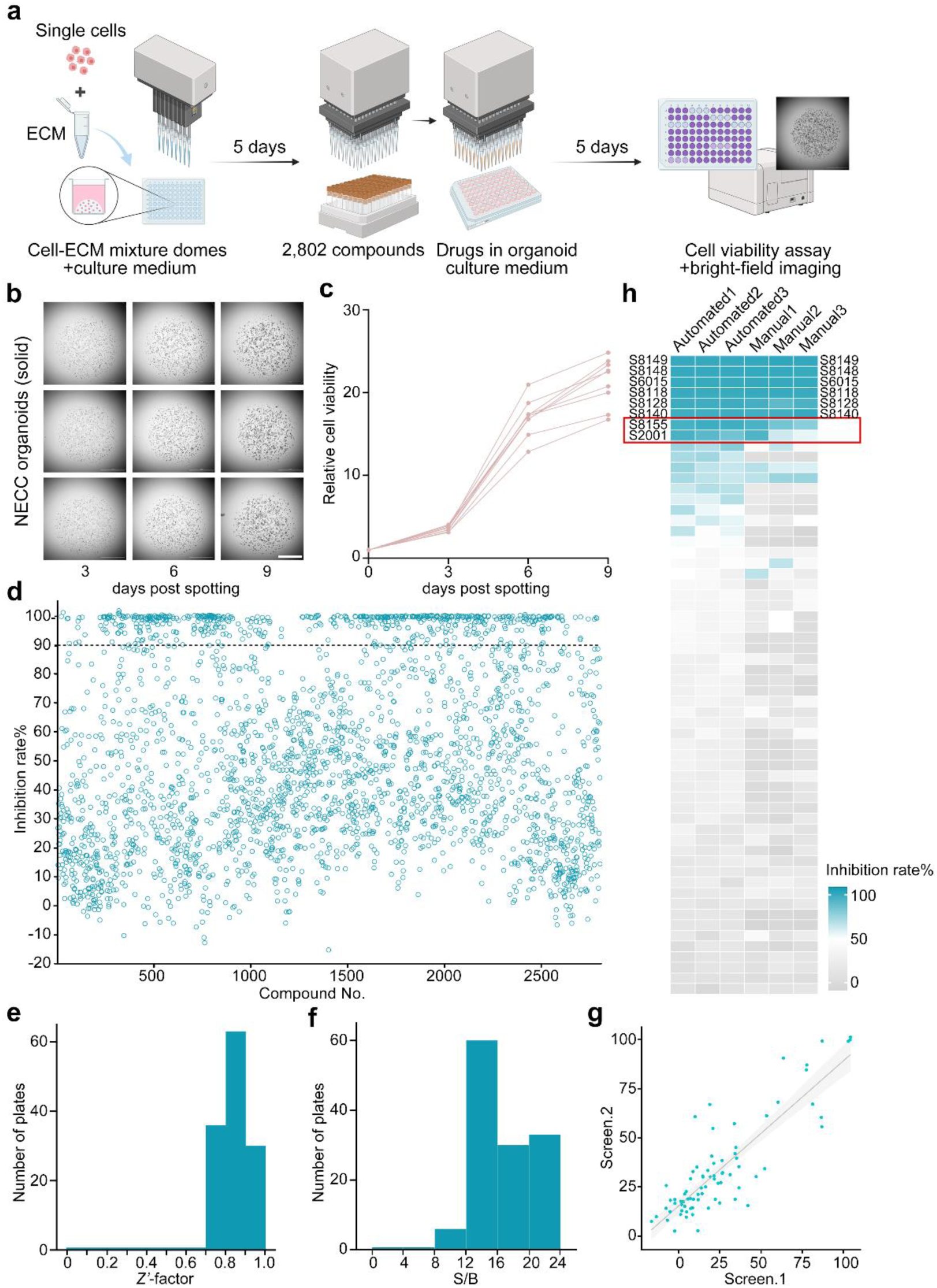
∣ The wp3D-OAHTS platform enables 2,802 drug screening on human NECC organoids. (**a**) Schematic workflow of conducting whole-process 3D NECC organoid drug screening using wp3D-OAHTS platform. The image was created with https://www.biorender.com/ (accessed on September 29, 2024). (**b**) Representative bright-field images showing NECC organoids growing after being automatically spotted. Scale bar, 1 mm. (**c**) Quantification of cell viability change ratio of NECC organoids (dots) after being automatically spotted. (**d**) Scatter plot of the inhibition rates of 2,802 compounds on NECC organoids, with 583 primary hits yielding over 90% inhibition rate tested at 10 μM. (**e**) Distribution of *Z′*-factors in 129 screening plates. (**f**) Distribution of signal-to-background (S/B) ratio parameters in 129 screening plates. (**g**) Correlation of the results between the different screen replicates using 77 randomly selected compounds. (**h**) Heatmap representation of inhibition rates for the randomly selected compounds used to compare automated screening with manual screening. Positive results with inhibition rates greater than 90% are annotated on both sides. Two additional positive results identified by automated screening compared to manual screening, S8155 (RSL3) and S2001 (Elvitegravir), are highlighted in red boxes.

In addition, we performed a traditional manual screening of a subset of these screening drugs. While the overall trends of inhibition rates obtained from automated and traditional manual screening were consistent, the automated screening identified two additional positive compounds not found in the manual screening: RSL3 (S8155) and Elvitegravir (S2001) (Fig. 3h). Importantly, bright-field images of organoids treated with either RSL3 or Elvitegravir revealed consistent, widespread cell death and organoid disintegration across all three replicate wells in the automated screening (Supplementary Fig. 2a, b). In contrast, manual screening exhibited significant differences among its replicates, suggesting that manual screening lacks the stability achieved by automated screening. These results demonstrate that automation, as a replacement for manual operation, is essential for conducting 3D organoid HTS, and our wp3D-OAHTS platform provides a robust tool for revealing differential compound effects, facilitating hit identification in automated 3D organoid HTS.

### Reverse drug concentration escalation identifies 7 top hits with potential for expanded indications in NECC

Among the 583 primary hits identified by our wp3D-OAHTS platform, more than half exerted lethal effects on NECC organoids (392 of 583), with inhibition rates ranging from 98% to 100% (Fig. 4a). This high efficacy suggested that 3D ECM-encapsulated organoid-based HTS could provide many potential therapeutic options for rare tumors. The majority of the primary hits (429 of 583) were investigational compounds that have been evaluated in clinical trials or are in preclinical development, while only 154 compounds were FDA-approved drugs. This fact suggests that there is a high probability that potential new NECC therapeutic candidates may emerge from compounds that have not yet received FDA approval (Fig. 4b). The 583 primary hits demonstrated a substantial degree of diversity and were categorized into 25 major classes based on their mechanisms of action (MOAs). Protein tyrosine kinase (PTK) inhibitors constituted the largest proportion at 12.18% (71 of 583), followed by cell cycle inhibitors and PI3K/Akt/mTOR inhibitors, each accounting for 8.40% (49 of 583) (Fig. 4c). These results indicate that high-throughput drug screening using 3D ECM-encapsulated organoids can reveal previously unidentified targets for rare diseases, providing a more diversified range of options for drug development.

**Figure 4.**
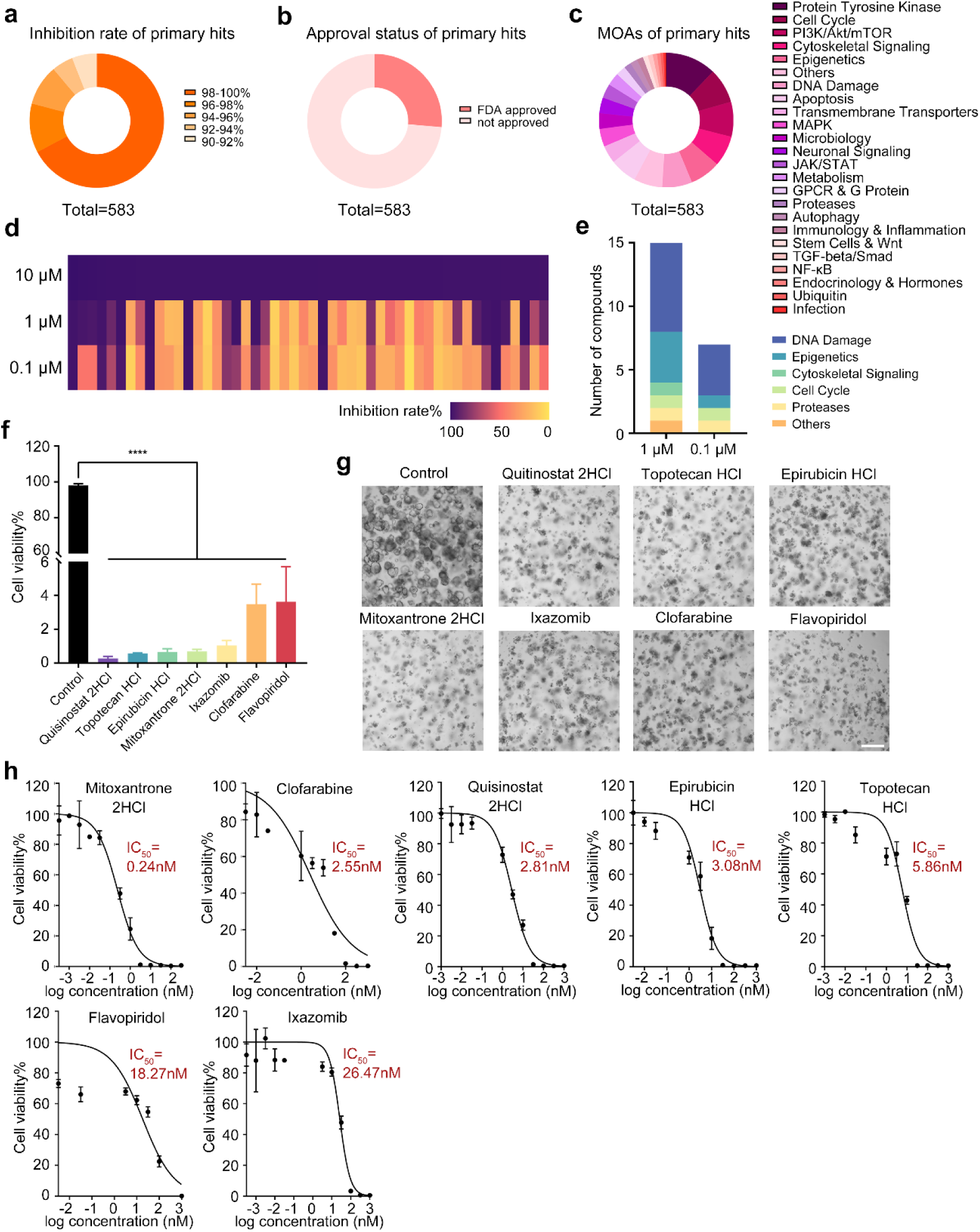
∣ Reverse drug concentration escalation identifies 7 top hits with potential for expanded indications in NECC. (**a-c**) Pie plots displaying the distribution of inhibition rates (a), approval status (b), and mechanisms of action (c) among the 583 primary hits. (**d**) Heatmap representation of inhibition rates for NECC organoids in secondary screening versus the 50 compounds selected from primary screening using concentrations of 1 μM and 0.1 μM. 15 of them showed >90% inhibition at 1 μM, and 7 of them were identified as top hits with >90% inhibition at 0.1 μM. (**e**) Bar plots showing the distribution of targeted pathways among the positive hits in secondary screening. (**f**) The normalized cell viability of organoids after treatment with the 7 top hits at 0.1 μM. Data are presented as mean ± SD (n=3). The p values were calculated by one-way ANOVA first and multiple comparisons test for further analysis. ****p < 0.0001. (**g**) Representative bright-field images of organoids in the control group and after treatment with the 7 top hits at 0.1 μM. Scale bar, 200μm. (**h**) Efficacy curves of the 7 top hits on NECC organoids. Data are presented as mean ± SD (n = 3).

To confirm and further narrow down the effective candidate drugs for NECC, we selected the top 50 compounds from our primary hits for a two-dose secondary screening (1 μM and 0.1 μM) against NECC organoids. Using serial dilution, we identified 15 and 7 candidates inducing growth inhibition >90% at 1 μM and 0.1 μM, respectively (Fig. 4d). The MOAs of the 15 candidates identified at the 1 μM working concentration included inhibitors of DNA damage (7 of 15), epigenetics (4 of 15), cytoskeletal signaling (1 of 15), cell cycle (1 of 15), proteases (1 of 15), and others (1 of 15). Upon reducing the working concentration to 0.1 μM, four DNA damage inhibitors, one epigenetic inhibitor, one cell cycle inhibitor, and one proteases inhibitor emerged as the top hits (Fig, 4e).

The 7 top hits, Quisinostat 2HCl, Topotecan HCl, Epirubicin HCl, Mitoxantrone 2HCl, Ixazomib, Clofarabine, and Flavopiridol, significantly reduced the viability of organoids and induced obvious cell death at 0.1uM (Fig. 4f, g). To better understand their potency, we generated full effective 12-point dose curves for each compound (Fig. 4h). From this assay, all of the 7 top hits were found to inhibit the growth of NECC organoids in a dose-dependent manner, with IC_50_ at the nanomolar level. Notably, five of these compounds (Mitoxantrone 2HCl, IC_50_ = 0.24 nM; Clofarabine, IC_50_ = 2.55 nM; Quisinostat 2HCl, IC_50_ = 2.81 nM; Epirubicin HCl, IC_50_ = 3.08 nM; and Topotecan HCl, IC_50_ = 5.86 nM) exhibited remarkably low IC_50_ below 10 nM, indicating their high potency and potential as promising therapeutic candidates for NECC, warranting further preclinical and clinical development.

Among the 7 top hits, Mitoxantrone 2HCl, Clofarabine, Epirubicin HCl, Topotecan HCl, and Ixazomib are FDA-approved drugs, while Quisinostat 2HCl and Flavopiridol are compounds previously being investigated in clinical trials (Supplementary Table 1). The current FDA-approved indications for these drugs do not include NECC, and the clinical trial data for Quisinostat 2HCl and Flavopiridol do not address NECC. These findings suggest their potential for expanding the therapeutic indications to include NECC.

### The representative drug Quisinostat 2HCl exhibits strong suppressive effects on NECC in vivo

To validate that our drug screening platform can serve as a novel tool for the discovery of potential therapeutics for clinical application, we focused on Quisinostat 2HCl, a non-FDA-approved compound with an IC_50_ of less than 10 nM. We examined the principal transcriptomic changes in NECC organoids after treatment with 0.1uM Quisinostat 2HCl for 48h and revealed a set of 3,815 differentially expressed genes (DEGs; 1,302 downregulated and 2,513 upregulated, absolute foldchange ≥ 2 and padj ≤ 0.05) (Supplementary Fig. 3a, b). Gene set enrichment analysis (GSEA) identified significant downregulation in multiple metabolic pathways such as cholesterol metabolism, steroid synthesis, and glycolysis/gluconeogenesis in Quisinostat 2HCl-treated organoids (Supplementary Fig. 3c-f). This finding suggests that Quisinostat 2HCl might inhibit the growth of NECC organoids by metabolism reprogramming.

To evaluate the anti-tumor effects of Quisinostat 2HCl in vivo, NECC organoids were transplanted subcutaneously in nude mice. When the transplanted tumors reached the designated volume, the mice were treated with 10 mg kg^-1^ Quisinostat 2HCl via intraperitoneal (IP) injection every other day for 20 days (Fig. 5a)^22^. The combination treatment of paclitaxel and carboplatin was included due to its established clinical application in the treatment of NECC and the favorable therapeutic effect observed in our NECC patient^23^. The growth of NECC xenografts in Quisinostat 2HCl-treated mice was significantly impaired compared with vehicle-treated mice, as evidenced by a marked decrease in both tumor volume and weight (Fig. 5b-d). It is notable that Quisinostat 2HCl treatment exhibited a higher inhibitory efficiency than the combination treatment of paclitaxel and carboplatin (Fig. 5b-d), which indicates its potential superiority as a monotherapy for NECC. These results collectively demonstrate that Quisinostat 2HCl possesses a potent capacity to inhibit the growth of NECC in vivo, highlighting its potential as a therapeutic agent for the treatment of NECC.

**Figure 5.**
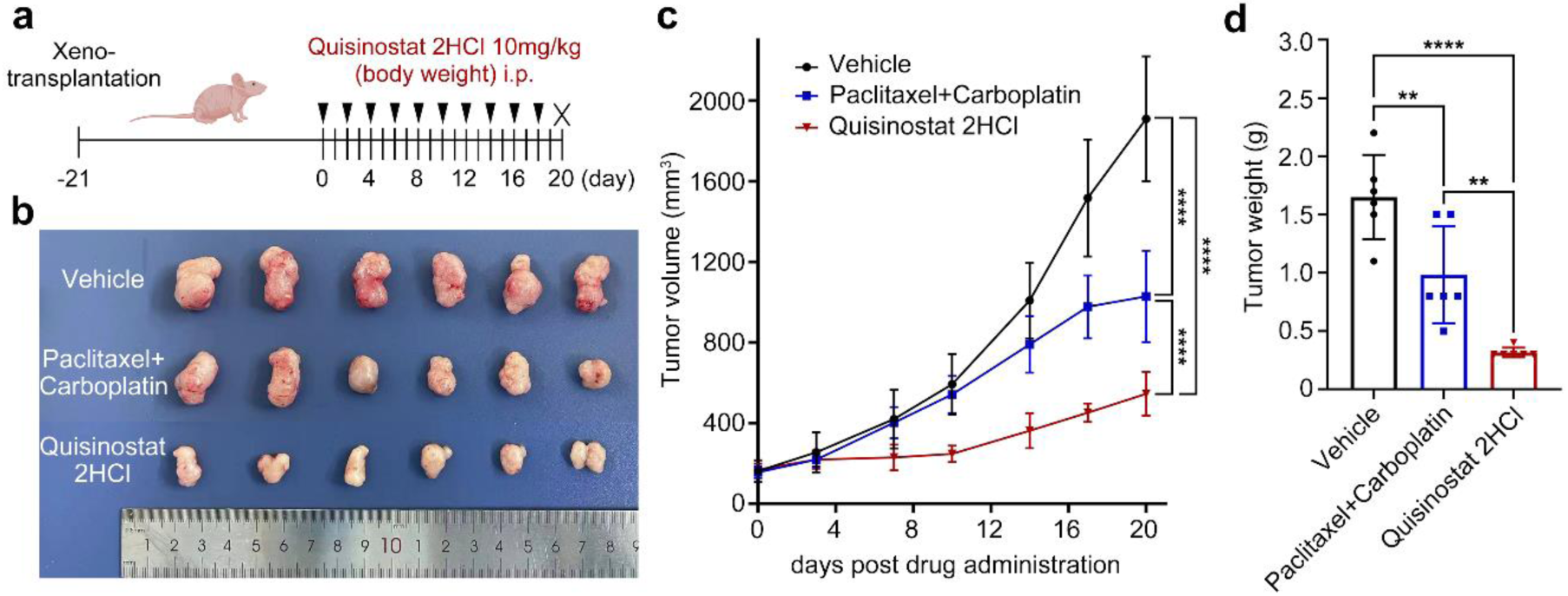
∣ The representative drug Quisinostat 2HCl exhibits strong suppressive effects on NECC in vivo. (**a**) Schematic diagram of the organoid xenograft model and in vivo drug treatment with Quisinostat 2HCl. Mice transplanted with NECC organoids received intraperitoneal injections of 10 mg kg^-1^ Quisinostat 2HCl every other day for 20 days. (**b**) Image of mice harboring NECC xenografts treated with Quisinostat 2HCl or Paclitaxel+Carboplatin or vehicle control. (**c**) Tumor growth curve of mice harboring NECC xenografts treated with Quisinostat 2HCl or Paclitaxel + Carboplatin or vehicle control. Data are presented as mean ± SD. N = 6 mice. (**d**) Tumor weight of mice harboring NECC xenografts treated with Quisinostat 2HCl or Paclitaxel + Carboplatin or vehicle control. Data are presented as mean ± SD. N = 6 mice. The p values were calculated by one-way ANOVA first and multiple comparisons test for further analysis. For tumor growth curve, the p values were calculated by two-way ANOVA. **p < 0.01; ***p < 0.001; and ****p < 0.0001.

### The wp3D-OAHTS platform permits precise drug responses by restricting false positives caused by organoid suspension

We have illustrated that even brief cultivation of organoids in a matrix-low suspension condition can induce transcriptomic alterations (Supplementary Fig. 1). To further investigate whether these changes in culture conditions influence the outcomes of organoid drug screening, we selected 6-7 compounds from each of the following inhibition rate ranges between 90-100%, 70-80%, 50-60%, and 30-40%, based on the initial 3D ECM-encapsulated organoid drug screening of 2,802 small molecules. These compounds were then tested against NECC organoids suspended in matrix-low conditions.

The conventional suspension-based organoids screening is similar to the whole-process 3D ECM-encapsulated organoids screening described here in that its initial phase involves encapsulating dissociated single cells in ECM, seeding them into wells of culture plates, and adding culture medium after gelation^11^. However, on the day of drug application, organoids in the suspension screening are detached from the ECM and resuspended in a culture medium containing 2% (vol/vol) ECM before being dispensed in the 96-well plates. In this setup, organoids are deprived of the supportive 3D environment and instead reside within a non-adherent suspension condition. Following 5 days of treatment with specific compounds, cell viability is assessed (Fig. 6a).

**Figure 6.**
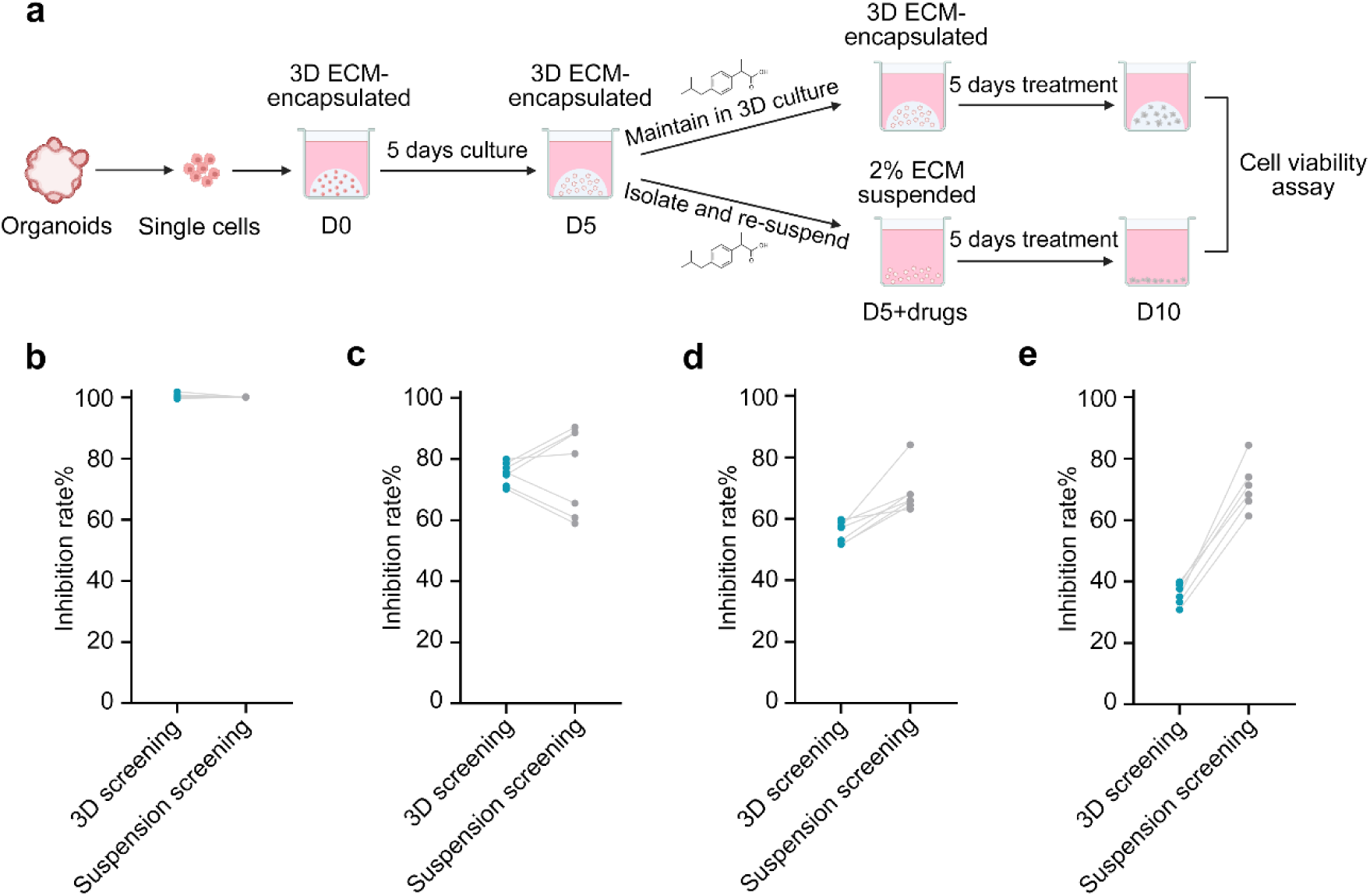
∣ The wp3D-OAHTS platform permits precise drug responses by restricting false positives caused by organoid suspension. (**a**) Schematic workflow of whole-process 3D ECM-encapsulated organoid-based screening and organoids screening suspended in matrix-low conditions. The image was created with https://www.biorender.com/ (accessed on September 29, 2024). (**b-e**) Comparative analysis of inhibition rates for 3D screening and suspension screening against selected compounds at 10 μM.

For small molecules exhibiting high sensitivity in NECC organoids, the outcomes between the whole-process 3D screening and suspension screening showed little discrepancy (Fig. 6b). Nevertheless, compounds that were previously shown to be non-responsive in the whole-process 3D screening exhibited enhanced sensitivity under suspension conditions (Fig. 6c-e). We found that over half of the small molecules with inhibition rates between 70-80% in 3D screening exhibited increased inhibition rates in suspension screening. Furthermore, all small molecules with inhibition rates between 50-60% in 3D screening showed increased inhibition rates in suspension screening. Meanwhile, those with inhibition rates between 30-40% in 3D screening also displayed inhibition rates with a more remarkable increase. This manifests that downgrading the 3D-cultured organoids to adapt to a 2D suspension system significantly increases the hit rate in organoid-based drug screening. However, these positive results may not accurately reflect the drug responses and resistance patterns of in vivo tumors, possibly attributed to the alterations in the 3D culture environment and transcriptomic landscape^24^. Employing wp3D-OAHTS platform for organoid drug screening enables the preservation of in vivo-like 3D structure of organoids, which is fundamental for the organoids to recapitulate drug responses of in vivo tumors, thereby fully leveraging the value of organoids in high-throughput drug screening.

## Discussion

Here we present a robust wp3D-OAHTS platform capable of automated spotting, organoid culturing, monitoring, and drug screening in 96-well plates. Organoids are maintained under 3D culture conditions encapsulated in an extracellular matrix component, which allows drug screening to be carried out in a state that most closely mimics the real tissues in vivo. Utilizing patient-derived 3D organoids for ex vivo drug screening has become a widely accepted practice. However, most HTS studies involving organoids are conducted in matrix-free or matrix-low culture medium due to challenges associated with automated HTS in ECM-encapsulated formats^1, 3, 11, 25, 26^. Specifically, there are two major challenges: (1) Firstly, the viscosity of extracellular matrix components is temperature-dependent. While the ECM remains liquidized at lower temperatures (2-8 °C), it solidifies as the temperature rises, making it difficult to accurately aspirate and dispense cell-matrix mixtures^12, 13^. (2) Secondly, organoid cultures require the cell-matrix mixture domes to be centrally located within the culture wells without adhering to the walls. HTS often employs high-density microplates such as 96-well and 384-well plates, where smaller well areas complicate precise domes positioning^27^. Additionally, it is necessary to shorten the dispensing time as much as possible and increase the screening output over a short period. To address these difficulties, researchers often choose to downgrade 3D organoid cultures to adapt to conventional simple HTS devices. In this study, we have shown that even shorter durations of culture in matrix-low conditions can lead to widespread alterations in the organoid transcriptome, leading to modifications in their drug responses (Supplementary Fig. 1, Fig. 6). Such strategies diminish the inherent advantages of organoids and make it difficult to reflect the true drug responses.

Previous studies have developed organoid seeding platforms that utilized microfluidic systems to create uniform cell-matrix droplets or micro-organospheres for HTS^28, 29^. These platforms generated organospheres using various separation oils, which introduced additional steps for droplet washing and oil removal prior to dispensing. Moreover, other studies have demonstrated the direct dispensing of liquefied cell-matrix mixtures onto the micropillar surface without introducing biphasic liquids^30^. This method replaces traditional culture plates with micropillar plates, achieving gelation, organoid culture, and drug exposure by inverting the micropillars into empty plates or plates containing medium or drugs. Our wp3D-OAHTS platform is designed to automatically spot cell-matrix mixtures into the center of wells in 96-well plates. Similar to the traditional manual organoid culturing process, this approach does not require the introduction of additional consumables or steps, thereby significantly simplifying the workflow and eliminating concerns about the culture state of the organoids.

The core module of our developed platform is a 3D ECM-encapsulated organoid spotter equipped with a cooling unit. The cooling unit maintains the cell-matrix mixtures at a low temperature within the sample plates. Prior to spotting, these mixtures are thoroughly mixed automatically and then aspirated by an 8-channel spotting head fitted with disposable tips, which spots the ECM droplets into the center of 96-well plate in a flying mode (0.1 seconds per well). After spotting, the domes are solidified on a heating unit, followed by the addition of culture medium via a 96-channel dispensing head. A robotic arm then transfers the culture plate into incubators. To facilitate continuous monitoring of organoid growth, an imaging and reading detector has been integrated into the automated system, allowing for the capture of images or viability assessments at predetermined time points, and providing real-time feedback on organoid growth. Based on the designed workflow for a 10-day screening cycle (where seeding, drug administration, and detection are each completed within a single day), our wp3D-OAHTS platform operating at maximum capacity can realize the screening of 8,694 compounds and the sceening of 43,730 compounds in only 14 days. This capability allows our platform to provide timely therapeutic guidance during the clinical treatment period, facilitating rapid decision-making based on PDOs.

We have utilized patient-derived 3D organoids from NECC, a rare malignancy with significant unmet clinical needs for novel drug development^14, 20, 21^, to perform HTS of 2,802 small molecules on our platform, to demonstrate the effectiveness, robustness, reproducibility, and stability of our screening system. We developed a two-step tiered HTS approach composed of primary screening and secondary screening, identifying 7 compounds that achieved over 90% inhibition rate of organoids at a working concentration of 0.1 μM, with 5 of them having an IC_50_ of less than 10 nM. This result suggests a potential clinical application in NECC treatment. Furthermore, we selected Quisinostat 2HCl, the only preclinical drug among these 5 candidates that has not yet received FDA approval, for in vivo efficacy validation. We found that Quisinostat 2HCl effectively inhibited the growth of NECC xenografts in mice. Quisinostat is an orally bioavailable potent pan-histone deacetylase inhibitor that has been tested in phase II clinical trials for cutaneous T-cell lymphoma and ovarian cancer but has not previously been reported to have any effect on neuroendocrine neoplasms^31, 32^. Our in vitro and in vivo results suggest that Quisinostat 2HCl has the great potential to be a therapeutic agent for neuroendocrine neoplasms treatment.

Compared to manual seeding of organoids in 96-well plates for drug screening, our automated system produces more consistent ECM domes across wells, contributing to a higher stability between replicate wells during drug screening. For example, in comparative testing of the manual and the automated systems, RSL3 and Elvitegravir were classified as positive hits by the automated screening but classified as negative hits by the manual screening. They accounted for 25% of the positive hits identified by the automated system. Based on bright-field imaging records, we observed significant variability between the replicate wells in the manual screening with inconsistent responses to drug treatment (Supplementary Fig. 2). In contrast, the automated screening demonstrated higher reproducibility across the three replicate wells. Adopting automated screening instead of manual screening can avoid the loss of 25% of positive results, which emphasizes the automation of HTS. Additionally, we compared the drug screening results of organoids cultured in suspension to those cultured in 3D conditions. We found that suspension cultures could increase organoid sensitivity to drugs, particularly for those that were insensitive under 3D culture conditions (Fig. 6). The finding may be attributed to the loss of 3D support in suspension culture, leading to a deviation from the in vivo tissue state and a shift towards a 2D-like condition. This possibly results in false positive outcomes similar to those obtained with 2D cell line screens and a high failure rate during clinical trials. Thus, compared to manual and suspension screening, our 3D ECM-encapsulated organoid-based automatic HTS demonstrates superior stability and efficacy, making it a definitive choice for organoid drug screening.

In summary, to maximize the advantages of 3D organoids in accurately reflecting in vivo tissue drug responses, we have developed an automated organoids HTS platform that operates organoids entirely in a 3D culture state. This platform achieves a more homogeneous, precise, and high-throughput seeding of 3D cell-matrix mixtures compared to the manual operation, which was previously challenging in HTS. Based on this, we have conducted a two-step tiered high-throughput drug screening of 2,802 small molecules using 3D ECM-encapsulated organoids for NECC, a rare malignancy with an urgent clinical need for new therapeutic options. We have identified seven candidate drugs with extremely low IC_50_, and we have validated the in vivo anti-tumor effect of one of these drugs, Quisinostat 2HCl. Furthermore, our wp3D-OAHTS platform allows for precise drug responses by restricting false positives caused by suspension cultures of organoids. These findings collectively demonstrate the superiority of that using 3D ECM-encapsulated organoids for high-throughput drug screening over the conventional manual system or the suspension culture-based organoid screening. The use of 3D ECM-encapsulated organoids would accelerate the discovery of new drugs for rare diseases.

## Methods

### wp3D-OAHTS platform design and assembly

The wp3D-OAHTS platform used for 3D ECM-encapsulated organoid-based HTS was designed and specified by our team at Fudan University, and subsequently customized and supplied by Novogene. This platform comprises four main modules: a 3D ECM-encapsulated organoids spotter, a liquid dispensing module, an automated incubating module, and a detecting module. The detailed sub-modules include a cooling unit, an 8-channel ECM spotting head, a heating-shaking unit, a 96-channel liquid dispensing head, a plate transfer unit, incubators, a robotic arm, a multimode reading unit, an imaging unit, a customized gripper unit, an automated plate storage unit, a workstation platform and a laminar flow hood.

### Organoid culture

In this study, the NECC patient provided written informed consent to participate in the clinical trial, in accordance with the principles of the Helsinki Declaration. The NECC biopsies and organoid establishment were conducted with the approval of the Medical Ethics Committee of Shanghai Chest Hospital (approval number: KS(Y)2078). The NECC, CC, GC, and CRC organoids were grown in drops of Organoid Culture ECM (bioGenous, M315066) in 24-well plates, overlaid with the corresponding medium. NECC organoids were maintained in Advanced DMEM/F-12 supplemented with penicillin–streptomycin (Invitrogen), HEPES buffer (10 mM, Gibco), GlutaMax (1X, Invitrogen), B27 supplement (1X, Invitrogen), N-acetyl-L-cysteine (1.25 mM, Sigma-Aldrich), Nicotinamide (5 mM, Sigma-Aldrich), A-83-01 (500 nM, bioGenous), SB202190 (500 nM, bioGenous), Y-27632 (5 µM, bioGenous), R-spondin1 (500 ng ml^−1^, bioGenous), Noggin (100 ng ml^−1^, bioGenous), EGF (50 ng ml^−1^, bioGenous), FGF10 (100 ng ml^−1^, bioGenous), and FGF7 (25 ng ml^−1^, bioGenous). Cervical Cancer Organoid Kit (bioGenous, K2169-CC) and Gastric Cancer Organoid Kit (bioGenous, K2179-GC) were used for the expansion of CC organoids and GC organoids, respectively. CRC organoids were maintained in Advanced DMEM/F-12 supplemented with penicillin–streptomycin (Invitrogen), GlutaMax (1X, Invitrogen), B27 supplement (1X, Invitrogen), N-acetyl-L-cysteine (1 mM, Sigma-Aldrich), Nicotinamide (10 mM, Sigma-Aldrich), A-83-01 (500 nM, bioGenous), SB202190 (3 µM, bioGenous), R-spondin1 (500 ng ml^−1^, bioGenous), Noggin (100 ng ml^−1^, bioGenous), EGF (50 ng ml^−1^, bioGenous), Gastrin-1 (100 µM, bioGenous), and PGE2 (1 nm, bioGenous). The medium was changed every 3-4 d.

### Organoid growth assessment

The growth of NECC, CC, GC, and CRC organoids after being automatically seeded by the wp3D-OAHTS platform was quantified by an increase in cell viability. Organoids grown in 20 μl ECM domes in 24-well plates were released from ECM in cold PBS, followed by centrifugation at 300 g for 3 min and resuspension in 1 ml of organoid dissociation solution (bioGenous, E238001). The organoids were digested at 37°C until they became single cells and were filtered using a 40-μm pore cell strainer. The filtrate was centrifuged at 300 g for 3 min. The cells were then mixed with culture medium and ECM (ratio=1:4) at the appropriate density and added to the wells of the sample plate, which was kept at 0 °C by the cooling unit. The cell-ECM mixtures (3 μl per well) were spotted into the centers of each well in a 96-well plate (Corning 3603 clear black flat-bottom microplates) using the 3D ECM-encapsulated organoids spotter. The plates were heated at 37 °C on the heating unit for 5 min for gelation. 100 μl of 1X LivingCell-Fluo™ Organoid Vitality Assay (bioGenous, E238004) reagent was added into each well and incubated in a 37 °C incubator for another 30 min. Fluorescence was measured at excitation 560 nm and emission 590 nm. Cell viability was detected and imaged every 3 days.

### Organoid drug screening on the wp3D-OAHTS platform

A compound library containing small molecular inhibitors and clinically used drugs was obtained from Selleck. NECC organoids grown in 20 μl ECM domes in 24-well plates were harvested, digested, and spotted into the 96-well plates as described above. Following gelation at 37 °C on the heating unit for 5 min, 100 μl of culture medium was dispensed into each well using a liquid dispensing head. The plates were then transferred to the detector for organoid imaging via a robotic arm before being returned to the incubators for further culture. Throughout the drug screening phase, each organoid plate was retrieved daily by the robotic arm from the incubators and transferred to the detector for imaging and monitoring, providing daily updates on the organoid growth status. After 5-day post-seeding of organoids, the automated liquid dispensing head performed the dosing procedure. Inhibitors were introduced into the culture medium at the final concentration of 10 μM during primary screening, while the 50 selected hits underwent secondary screening at final concentrations of 1 μM and 0.1 μM. For dose curve confirmation, the 7 top hits were added to the culture medium at different final concentrations. Positive and negative controls were also included. After 5 days of treatment, the plates were retrieved by the robotic arm and imaged using the detector, followed by placement in the liquid dispensing module for medium removal. 100 μl of 1x LivingCell-Fluo™ Organoid Vitality Assay (bioGenous, E238004) reagent was added into each well and incubated in a 37 °C incubator for another 30 min. Fluorescence was measured at excitation 560 nm and emission 590 nm. To determine the percentage reduction in treated versus control organoids in cytotoxicity assays, the following formula was used:

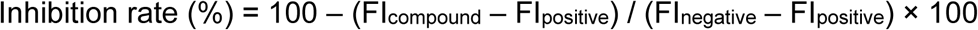

Where: FI = Fluorescence Intensity.

To calculate IC_50_, viability was normalized relative to the average of positive controls and the highest concentration drug-treated conditions. Efficacy curves were generated using Prism GraphPad.

### Drug validation in xenograft model

6-week BALB/c-Nude mice (Strain NO. D000521) were purchased from GemPharmatech (Nanjing, China). All animal experiments were approved by the Institutional Animal Care and compliant with all relevant ethical regulations regarding animal research. Organoids were transplanted subcutaneously with 100,000 cells in 100 μl 50% ECM per injection in PBS. Tumor volume was measured with calipers. After the tumor volume of each mouse reached 150-200 mm^3^, the mice were randomly divided into 3 groups for treatment with drugs or vehicles. Quisinostat 2HCl was administrated at 10 mg kg^-1^ via intraperitoneal injection every other day. Combination therapy of paclitaxel (10 mg kg^-1^) with carboplatin (50 mg kg^-1^) was administrated via intraperitoneal injection once a week. Mice were sacrificed 20 days after drug administration. The animal ethics (2020JS038) was approved by Laboratory Animal Center, School of Life Sciences, Fudan University.

### RNA-seq

Total RNA of NECC organoids suspended in culture medium containing 2% (vol/vol) ECM, NECC organoids encapsulated in 3D ECM, NECC organoids treated with 0.1 μM Quisinostat 2HCl or PBS for 48 h were isolated using the RNAprep Pure Micro Kit (Tiangen Biotech) and then sent for sequencing (Novogene). High-throughput sequencing was performed using Illumina NovaSeq X Plus for 3 biological replicates, respectively. The raw sequencing data quality was checked by FastQC. HISAT2 was then used to align clean reads to the human reference genome (GRCh38) with default parameters. Bam files were sorted by Samtools and count matrices were generated by StringTie.

Downstream analysis was processed in R v4.2.1. Briefly, DESeq2 v1.38.3 was used to identify DEGs. For comparison between NECC organoids suspended in culture medium containing 2% ECM and encapsulated in 3D ECM, genes with padj ≤ 0.05 and absolute foldchange ≥1.5 were regarded as DEGs. For comparison between NECC organoids treated with 0.1 μM Quisinostat 2HCl or PBS, genes with padj ≤ 0.05 and absolute foldchange ≥2 were regarded as DEGs. ClusterProfiler v4.6.2 was used to do KEGG enrichment and GSEA, and gene expression heatmaps were generated by pheatmap v 1.0.12.

### Statistical analysis

We employed one-way ANOVA test and two-way ANOVA test to analyze the experimental results. Analyses were conducted on GraphPad Prism 8 statistical software. All values are represented as means ± SD. The value of \**P* < 0.05, \*\**P* < 0.01, \*\*\**P* < 0.001, \*\*\*\**P* < 0.0001 was considered significant.

## Data Availability

The data used and/or analyzed during the current study are available from the corresponding author on request.

## Acknowledgements

This work was supported by grants from the National Natural Science Foundation of China (82372663), the Key Research and Development Program of Yunnan Province (202302AA310024), the Key Research and Development Program of Jiangxi Province (20232BBG70024), the Natural Science Foundation of Shandong Province (ZR2023LSW008).

## Author contributions

Z.X. and B.Z. conceived the study; Z.X. performed the experiments; Z.X., H.Y., Y.Z. and E.E.D analyzed the data; B.Z. supervised the work; and Z.X. and B.Z. wrote the manuscript.

