## Supplementary Figure 1-3 for "Whole-Process 3D ECM-Encapsulated Organoid-Based Automated High-Throughput Screening Platform Accelerates Drug Discovery for Rare Diseases"

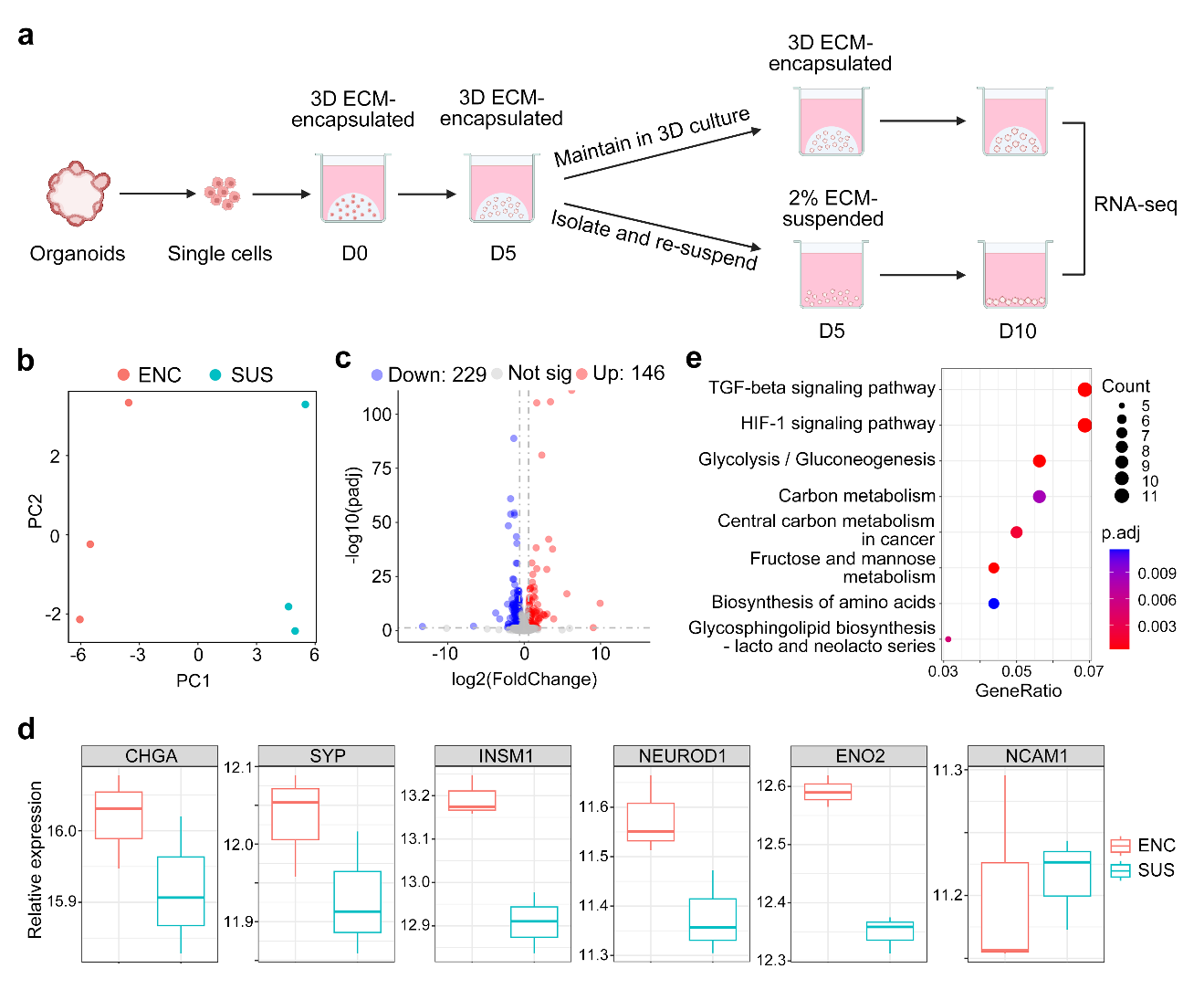

**Supplementary figure 1 ∣ Traditional organoid drug screening suspended in matrix-low conditions induces transcriptomic modifications.** (**a**) Schematic workflow of RNA-seq for whole-process 3D ECM-encapsulated organoid cultures and traditional suspended organoid cultures. The image was created with https://www .biorender.com/ (accessed on September 29, 2024). (**b**) Principal component analysis of encapsulated (n=3) and suspended (n=3) organoids. ENC, encapsulated organoids. SUS, suspended organoids. PC, principal component. (**c**) Volcano plot of differentially expressed genes (DEGs) in suspended versus encapsulated organoids. Genes with padj ≤ 0.05 and absolute foldchange ≥1.5 were regarded as DEGs. (**d**) Boxplot of the relative expression of tumor markers in encapsulated and suspended organoids. (**e**) KEGG analysis of DEGs in suspended organoids, compared to encapsulated organoids.

**Supplementary figure 2**

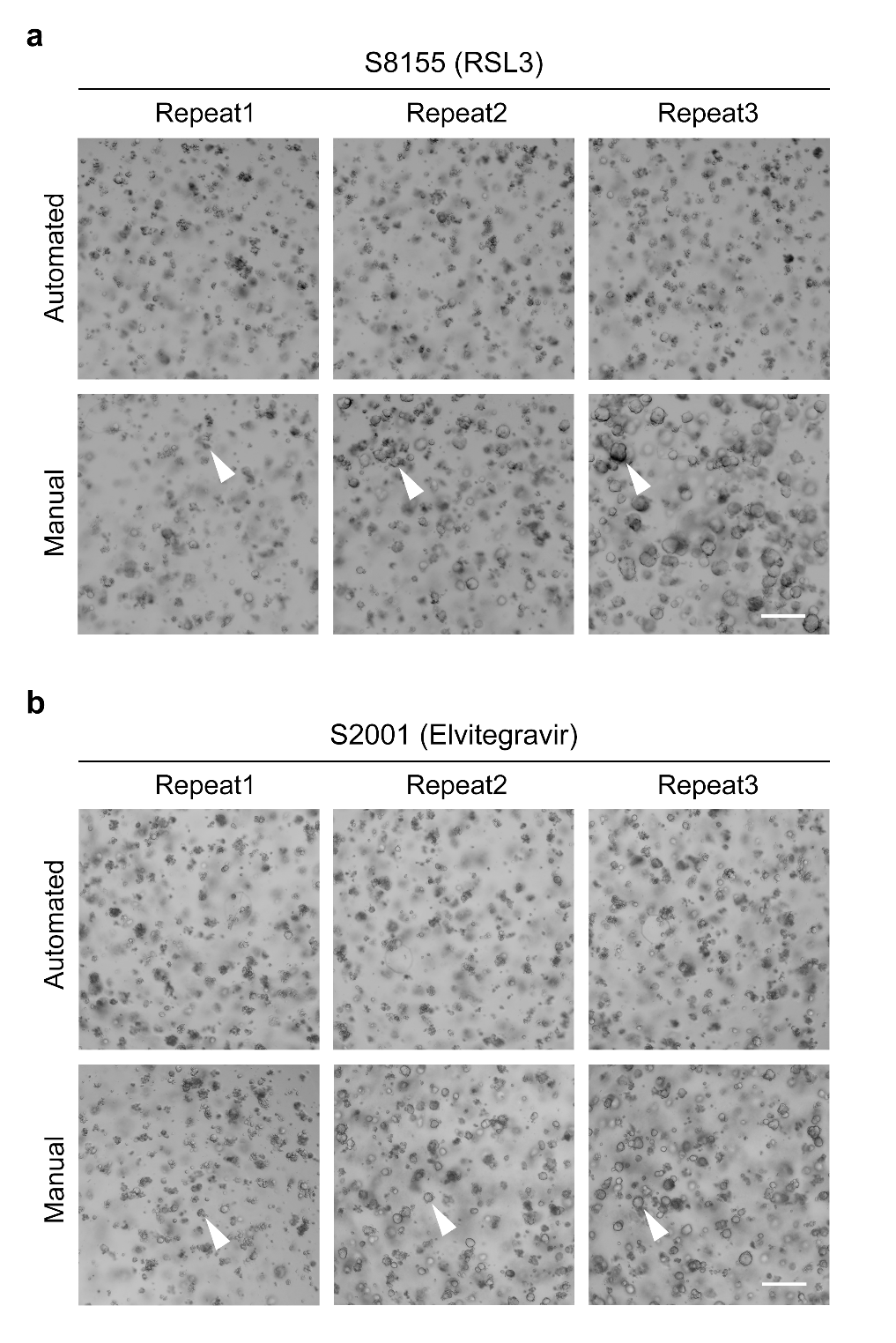

**Supplementary figure 2 ∣ Automated 3D screening exhibits higher stability compared to manual 3D screening and recovers positive hits missed by manual operation.** (**a**) Bright-field images of organoids in three replicate wells for automated (top) and manual (bottom) screening after treatment with S8115 (RSL3) at 10 μM. Scale bar, 200μm. White arrows indicate the differences in organoid growth in the three replicate wells of the manual screening. (**b**) Bright-field images of organoids in three replicate wells for automated (top) and manual (bottom) screening after treatment with S2001 (Elvitegravir) at 10 μM. Scale bar, 200μm. White arrows indicate the differences in organoid growth in the three replicate wells of the manual screening.

**Supplementary figure 3**

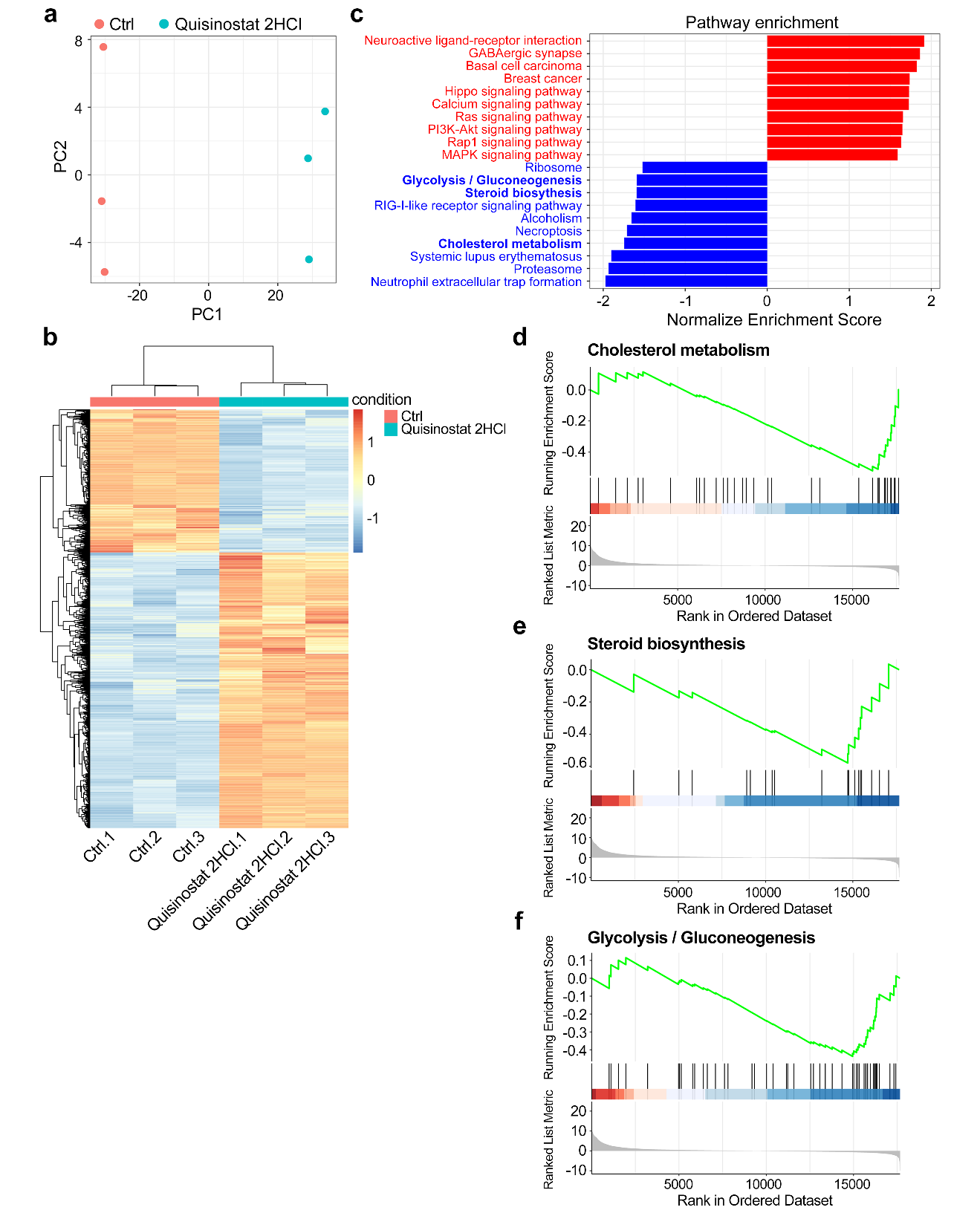

**Supplementary figure 3 ∣ Quisinostat 2HCl might inhibit the growth of NECC organoids by metabolism reprogramming.** (**a**) Principal component analysis of control (n=3) and Quisinostat 2HCl-treated (n=3) organoids. PC, principal component. (**b**) mRNA expression heatmap of differentially expressed genes for control (n=3) and Quisinostat 2HCl-treated (n=3) organoids. (**c**) Gene set enrichment analysis (GSEA) of Quisinostat 2HCl-treated versus control organoids. Upregulated pathways are shown in red text, and downregulated pathways are shown in blue text. (**d-f**) GSEA analysis showing enrichment of cholesterol metabolism, steroid biosynthesis, and glycolysis/gluconeogenesis pathway in Quisinostat 2HCl-treated versus control organoids.
